# Conserved interhemispheric morphogenesis in amniotes preceded the evolution of the corpus callosum

**DOI:** 10.1101/2024.12.03.625459

**Authors:** Ryota Noji, Mari Kaneko, Takaya Abe, Hiroshi Kiyonari, Yukihiro Nishikawa, Takuma Kumamoto, Hitoshi Gotoh, Chiaki Ohtaka-Maruyama, Katsuhiko Ono, Tatsuya Yoshizawa, Tadashi Nomura

## Abstract

The corpus callosum (CC) is the large axon bundle connecting the telencephalic hemispheres. The CC is formed exclusively in placental mammals, and the lack of comparable structures in other amniotes obscures the evolutionary origin of the CC. We here demonstrate that interhemispheric remodeling, a prior developmental step for CC formation, is highly conserved in non-mammalian amniotes, such as reptiles and birds. In these animal groups, the spatiotemporal dynamics of interhemispheric remodeling are tightly connected with distinct commissural formations. We observed a high degree of similarity between the mammalian CC and reptilian rostral pallial commissure, (RPC) and significant modifications in the avian pallial projection. Furthermore, we determined that Satb2 plays crucial roles in interhemispheric remodeling, which is associated with proper formation of both the CC and RPC in mice and geckoes, via the use of CRISPR-mediated gene-targeting. Our findings suggest that developmental mechanisms for midline remodeling were already present in the common ancestor of amniotes, which contributed to the evolution of eutherian-specific CC formation.

## INTRODUCTION

The corpus callosum (CC) is the prominent axon bundle connecting the two cerebral hemispheres and plays crucial roles in the integration of interhemispheric information processes. The CC is formed exclusively in placental (eutherian) mammals, and comparable anatomical structures do not exist in other extant mammalian taxa (marsupials and monotremes) or in non-mammalian amniotes, such as reptiles and birds^1,2^. Notably, in these animal groups the two cerebral hemispheres are interconnected with the anterior commissure and the hippocampal commissure^3^, which are highly conserved commissural pathways in all amniote species^2,4^. However, it remains unclear how the new route of interhemispheric connections appeared evolved in eutherian lineages.

Several studies have demonstrated multiple developmental steps for mammalian CC formation^2,5–7^. In early stages of mammalian brain development, the two cerebral hemispheres are separated by interhemispheric fissure (IHF), which is composed of leptomeningeal cells and an extracellular matrix^8^. As development progresses, specific astroglial cells in the telencephalic midline, such as midline zipper glia (MZGs), indusium griseum glia, and glial wedges, play essential roles in the retraction of the IHF^9,10^, through the degradation of leptomeningeal cells and the intercalation of midline tissue^11,12^. The remodeling process of the IHF is crucial for CC formation because it involves bridging the two hemispheres before pioneer callosal axons reach the midline^12–15^. Defects in interhemispheric remodeling are strongly associated with congenital agenesis of the CC in humans^7,16^. However, how the dynamic process of midline remodeling prior to CC development evolved in ancestral mammals remains unclear. Consistent with the lack of CC formation, IHF retraction is not detected in marsupials^11^, while the presence of midline zipper glia in these animal groups obscures the evolutionary origin of IHF remodeling and the underlying molecular mechanisms.

Special AT-rich sequence-binding protein 2 (Satb2) plays crucial roles in mammalian neocortical development. In the developing murine neocortex, Satb2 regulates the differentiation of upper cortical neurons that contribute to CC bundle formation^17–20^. The loss of Satb2 function results in the fate conversion of upper cortical neurons to deep layer neurons, whereby callosal axons misroute to subcortical regions^17–19,21^. Furthermore, the differential timing of Satb2 expression in the developing neocortex provides a route of species-specific commissural connections, suggesting that heterochronic expression of Satb2 contributes to eutherian and marsupial-specific interhemispheric projections^22^. Notably, several studies implicated that agenesis of CC in Satb2-deficient mice is also associated with defects in midline fusion^23^. However, the functional contribution of Satb2 to the midline glial has not been examined in detail.

Here we report a remarkable degree of conservation in the interhemispheric remodeling process in non-mammalian amniotes, geckoes, turtles, and birds, despite the lack of a CC in these animal groups. By performing a detailed histological analysis, we observed that distinct developmental mechanisms confer rostral and caudal IHF remodeling, which is tightly associated with the route of other commissural pathways: the rostral pallial commissure (RPC) and HC. Furthermore, using CRISPR-mediated gene targeting, we observed that Satb2 plays crucial roles in interhemispheric remodeling, which is associated with proper formation of both the CC and RPC in mice and geckoes. Our findings suggest that developmental mechanisms for interhemispheric remodeling were already present in the common ancestor of amniotes, which contributed to the evolution of eutherian-specific CC formation.

## RESULTS

### Interhemispheric remodeling is highly conserved in the developing amniote brain

To address the evolutionary conservation of the interhemispheric remodeling process in non-mammalian brains, we first compared midline morphogenesis of the telencephalon in various amniote species, including mice (*Mus musculus*), Chinese softshell turtles (*Pelodiscus sinensis*), domestic chicks (*Gallus gallus*), and Madagascar ground geckoes (*Paroedura picta*). Hematoxylin and Eosin (H&E) staining of serial horizontal sections throughout development confirmed that in the early embryonic stages of all the examined species, telencephalic hemispheres are connected only by thin epithelial tissue (telencephalic hinge), whereas the rest of the midline region is completely separated by the IHF (Figure 1A-D). In mice, after E14, the IHF is progressively retracted and both sides of the septal tissues are fused to bridge the telencephalic hemispheres (Figure 1A). Notably, these interhemispheric remodeling processes (i.e., IHF retraction and septal fusion) were also detected in turtles, chicks, and geckoes, as telencephalic development progressed (Figure 1B-D). Exceptionally, in chicken telencephalon the extent of IHF retraction was limited to the caudal part of the septum, and the rest part of the septal midline remained to be separated (Figure 1C). In mice, interhemispheric remodeling occurs from the caudal-ventral to the rostral-dorsal direction during development^12^. The ratio of the IHF length to total midline length revealed that stage-dependent IHF retraction occurs progressively in ventral to dorsal direction in both the mouse and the gecko telencephalon (Figure 1E-G), suggesting that the spatial and temporal progression of interhemispheric remodeling is highly conserved in amniotes.

**Figure 1.**
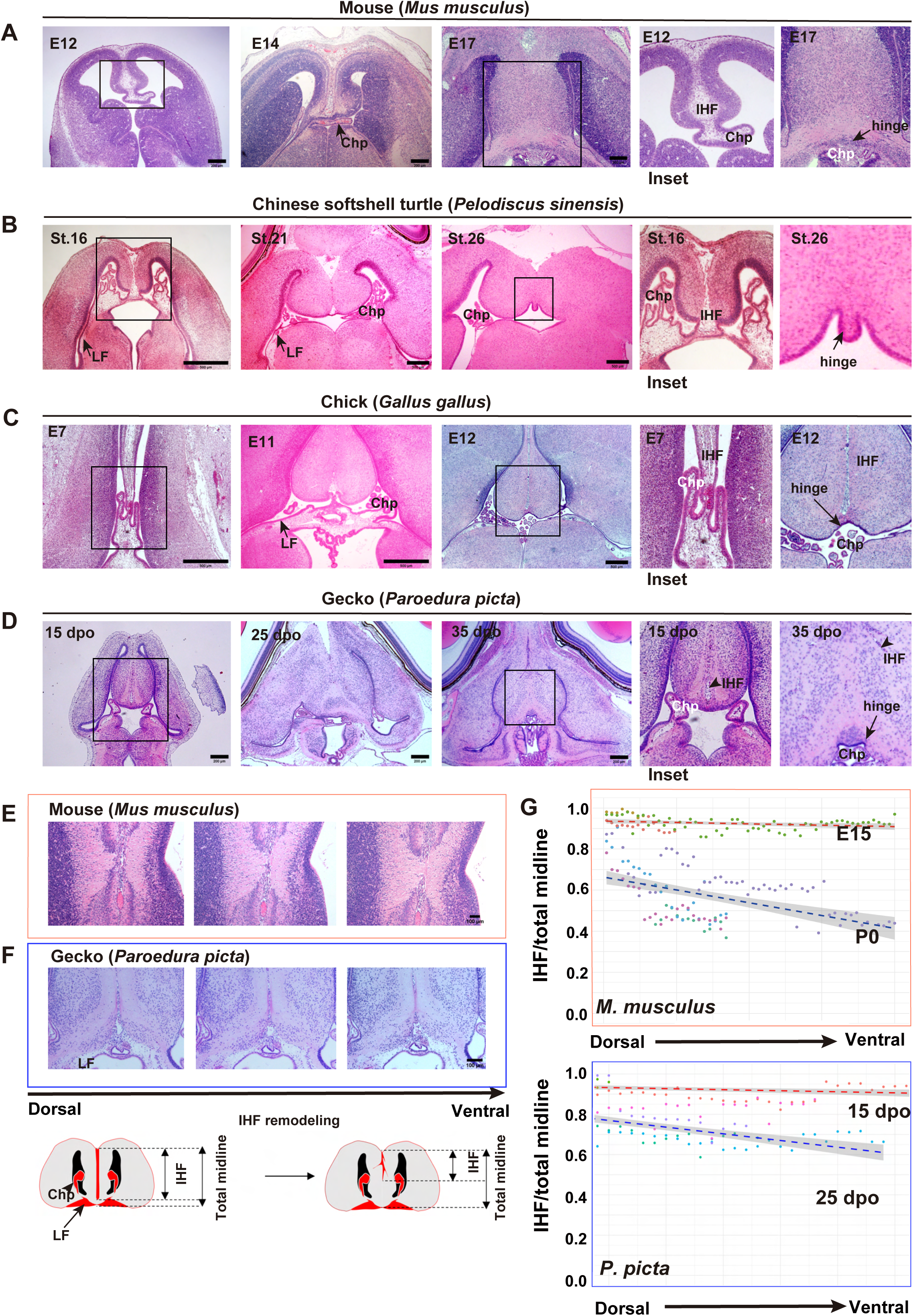
Interhemispheric remodeling is common in the developing amniote brains. (A-D) Horizontal sections of developing mouse (A), Chinese softshell turtle (St. 16 and St. 26; B), chick (C), and Madagascar ground gecko (D) forebrains. All sections were stained with hematoxylin and eosin. (E-G) Progressive regression of interhemispheric fissure (IHF) from the dorsal to ventral planes in the mouse (E) and gecko (F) forebrains. (G) Quantitative analysis of the IHF regression in a mouse (upper panel) and gecko (lower panel) at different embryonic stages (mouse: E15, N=3 animals and P0 N=4 animals; gecko: 15 dpo N=3 animals and 25 dpo N=4 animals). The formula for simple linear regression to various groups (mouse: E15, y=0.9359841−0.0002687x: R^2^=0.06353; P0, y=0.6630286−0.0024960x, R^2^=0.2498; Geckos:15dpo, y=0.9430885−0.0010498x, R^2^=0.1184; 25dpo, y=0.8253532−0.0045618x, R^2^=0.1938). Horizontal axis: IHF/total IHF, Vertical axis: anatomical positions. The graph shows the ratio of IHF length to total midline length along the dorsal-ventral axis of the forebrain based on anatomical landmarks. For each individual, a dot represents a section and is color coded; early developmental stages represent warm colors, while late developmental stages represent cold colors. IHF : Interhemispheric fissure. Chp : Choroid plexus. LF: Longitudinal Fissure.

### MZG-dependent degradation of leptomeningeal components occurs in reptiles

To address whether the interhemispheric remodeling in non-mammalian amniotes is dependent on midline glial activities, we next examined the expression of several glial markers in the reptilian and avian midline regions. We detected the accumulation of GFAP-positive cells and phosphorylated vimentin (Ser55) -positive mitotic radial glia^24^ in the IHF and telencephalic hinge in developing turtles, geckoes and chicks (Figure 2A), suggesting that interhemispheric remodeling in non-mammalian species is mediated by specific glial components, such as the MZG and glial wedge, as in the case of eutherian mammals. It has been shown that MZG-dependent degradation of leptomeninges is crucial for the retraction of the IHF in the developing mouse brain^12^. We detected the degenaration of laminin-positive leptomeninges and its degradation during astroglial interculation(GFAP-positive glial fibers) in the developing gecko (Figure 2B). In addition, it was suggested that astroglial interculation occurs in conjunction with the aforementioned regression of the IHF from st.21 to st.26 in the developing turtle(Figure 2C). Furthermore, we corroborated the activity of pan-matrix metalloproteinase (MMP) in the laminin-positive leptomeninges adjacent to GFAP-positive glial fibers in the developing turtle midline region (Figure2D, Supplementary Movie1), suggesting that MMP-mediated degradation of leptomeningeal components occurs during interhemispheric remodeling in reptiles, as reported in mice^12^.

**Figure 2.**
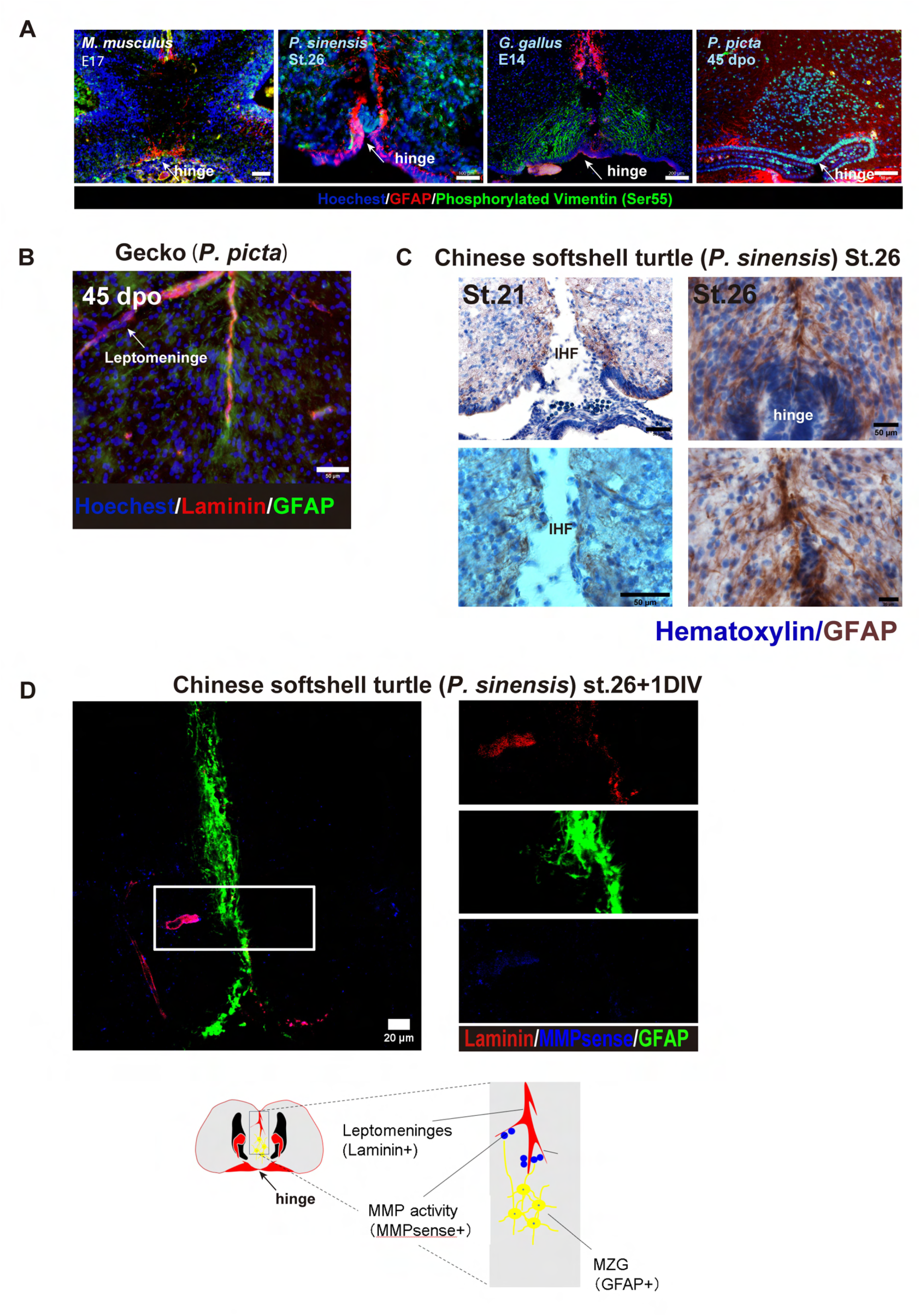
MZG-dependent degradation of leptomeningeal cells in non-mammalian amniotes. (A) GFAP-positive or phosphorylated vimentin-positive midline glial cell components in the developing mouse, turtle, chick, and gecko forebrains. (B) Distribution of laminin-positive leptomeninges and GFAP-positive midline zipper glia (MZG)-like fibers in the gecko forebrain. (C) GFAP-positive fibers before IHF regression (left) and during IHF regression (right) in the turtle forebrain by immunohistochemistry. The lower panels show high magnification of each image. (D) Detection of pan-matrix metalloproteinase (MMP) activity during IHF remodeling of the turtle forebrain in slice culture(DIV: Day In Vitro) and schematic drawings. IHF : Interhemispheric fissure.

### Epithelial remodeling occurs during caudal-ventral interhemispheric remodeling

As observed histologically, telencephalic hemispheres are joined with the lamina terminalis, which gives rise to the choroid plexus during embryogenesis. We confirmed that at early embryonic stages the two hemispheres are connected with the primordial choroid plexus in all examined amniote species (Figure 1A-D). However, at later stages the choroid plexus is disconnected from the telencephalic hinge that bridges the caudal-ventral part of the septum. We examined serial oblique sections of the septum from the dorsal to ventral planes, which represent the temporal process of interhemispheric remodeling, and corroborated the gradual disconnection of the choroid plexus from the hinge tissue in the mouse, turtle, chick, and gecko midlines (Figure 3A-E). We also confirmed the accumulation and invasion of phosphorylated vimentin-positive glial fibers into the lamina terminalis, which coincided with the disruption of the laminin-positive leptomeninges, during the remodeling process in chick embryos (Figure 3F, G, Supplementary Movie2). These lines of evidence suggest that dynamic rearrangement of epithelial tissues occurs at the caudal-ventral part of the septum. In the developing chicken brain, the distributions of vimentin-positive fibers and GFAP-positive MZG-like cells were clearly separated in the interhemispheric midline, which were coincided with remodeling (caudal) and non-remodeling (rostral) regions of the interhemispheric fissure (Figure 2A). This finding suggests that caudal and rostral interhemispheric remodeling are mediated by distinct glial components, and despite the presence of MZG-like cells, IHF retraction does not occur in the rostral part of the septum in chicken brains.

**Figure 3.**
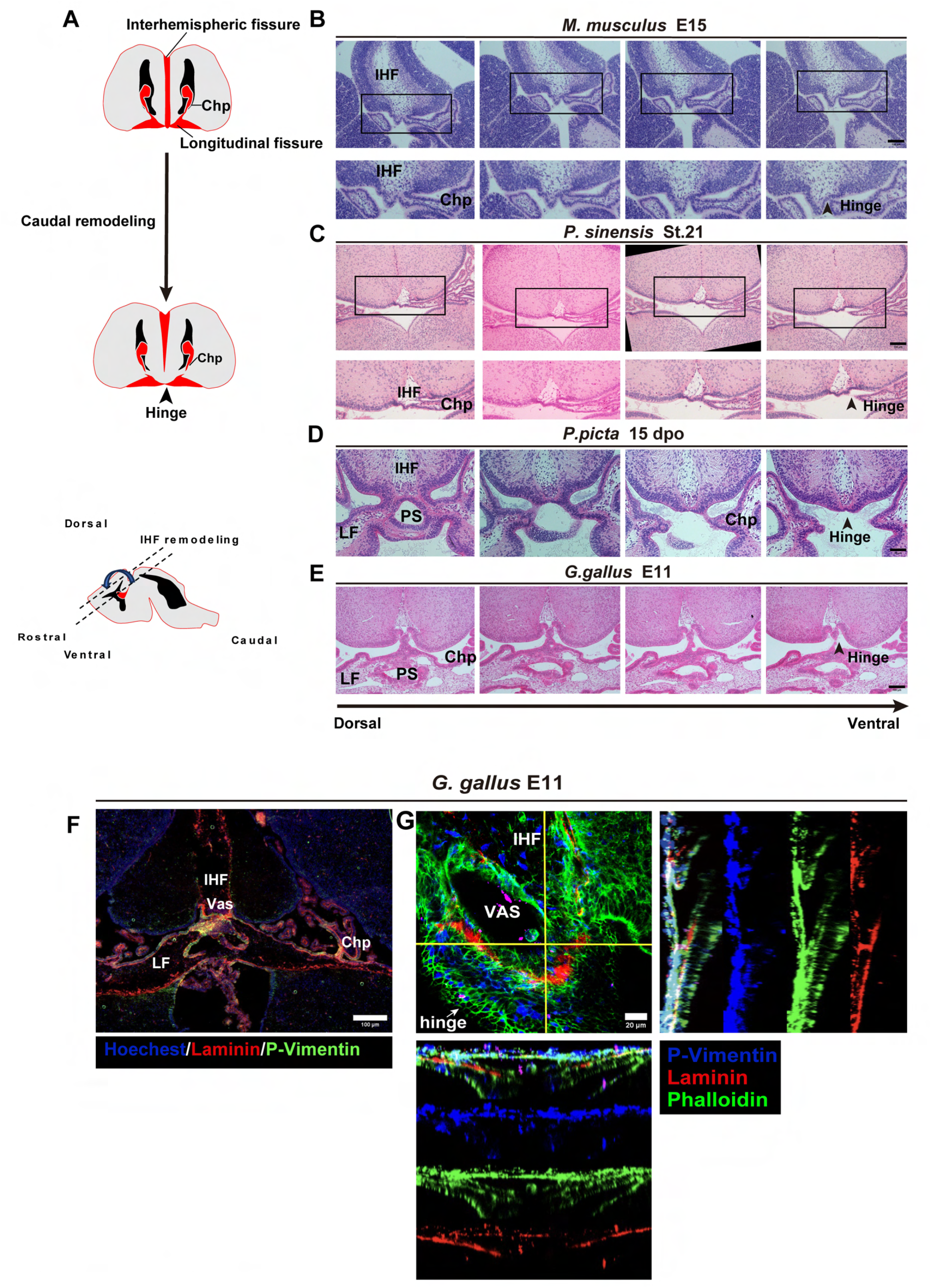
Epithelial remodeling during the caudal-ventral interhemispheric remodeling. (A) Schematic drawings of caudal remodeling procedures. (B) Serial sections of the dorsal to ventral plane demonstrating epithelial remodeling of hinge regions in the developing mouse (B), turtle (C), gecko (D), and chick (E) forebrains. (F) Distribution of laminin and phosphorylated vimentin at the lamina terminalis of chick forebrain. (G) A confocal image showing invasion of phosphorylated vimentin-positive glia into the leptomeninges in the chick forebrain. Histological images of turtles (C), geckos (D), and chicks (E) were obtained from the same individuals as in Figure 1. IHF : Interhemispheric fissure. Chp : Choroid plexus. LF: Longitudinal Fissure. Vas : Vasculature.

### Rostral and caudal remodeling bridge the pathway of distinct pallial commissures

To determine whether interhemispheric remodeling in non-mammalian amniotes provides substrates for distinct commissural pathways, we traced commissural axons in gecko hemispheres and compared them with those in murine brains. Unilateral deposition of a lipophilic tracer on the gecko dorsal cortex resulted in retrograde labeling of cells in the contralateral cortex (Figure 4B), as in the case of the labeling of callosal projections in the mammalian neocortex (Figure 4A). Thus, the reptilian telencephalic hemispheres are connected with commissural axons at positions similar to those of the mammalian CC. To compare topographic patterns of commissural pathways in mice and geckoes, we combined immunostaining with an anti-GAP43 antibody, a conserved marker for commissural axons, and anterograde axon tracing. We confirmed that distinct routes of commissural axons that connect the rostral and caudal parts of gecko hemispheres, which correspond to the rostral and caudal pallial commissure (RPC and CPC), respectively (Figure 4C). Notably, the positions of RPC and CPC in geckoes is coincided well with the anatomical positions of the CC and HC in mice (Figure 4C). Furthermore, we found that GAP43-positive axons do not cross the midline before IHF remodeling in the developing chicken and turtle brains (Figure S1). These data suggest that rostral and caudal interhemispheric remodeling play crucial roles in the routing of distinct commissural pathways; in particular, rostral midline remodeling provides substrates for both the CC and RPC, whereas the caudal remodeling bridges the route of the HC in both the mammalian and reptilian brain.

**Figure 4.**
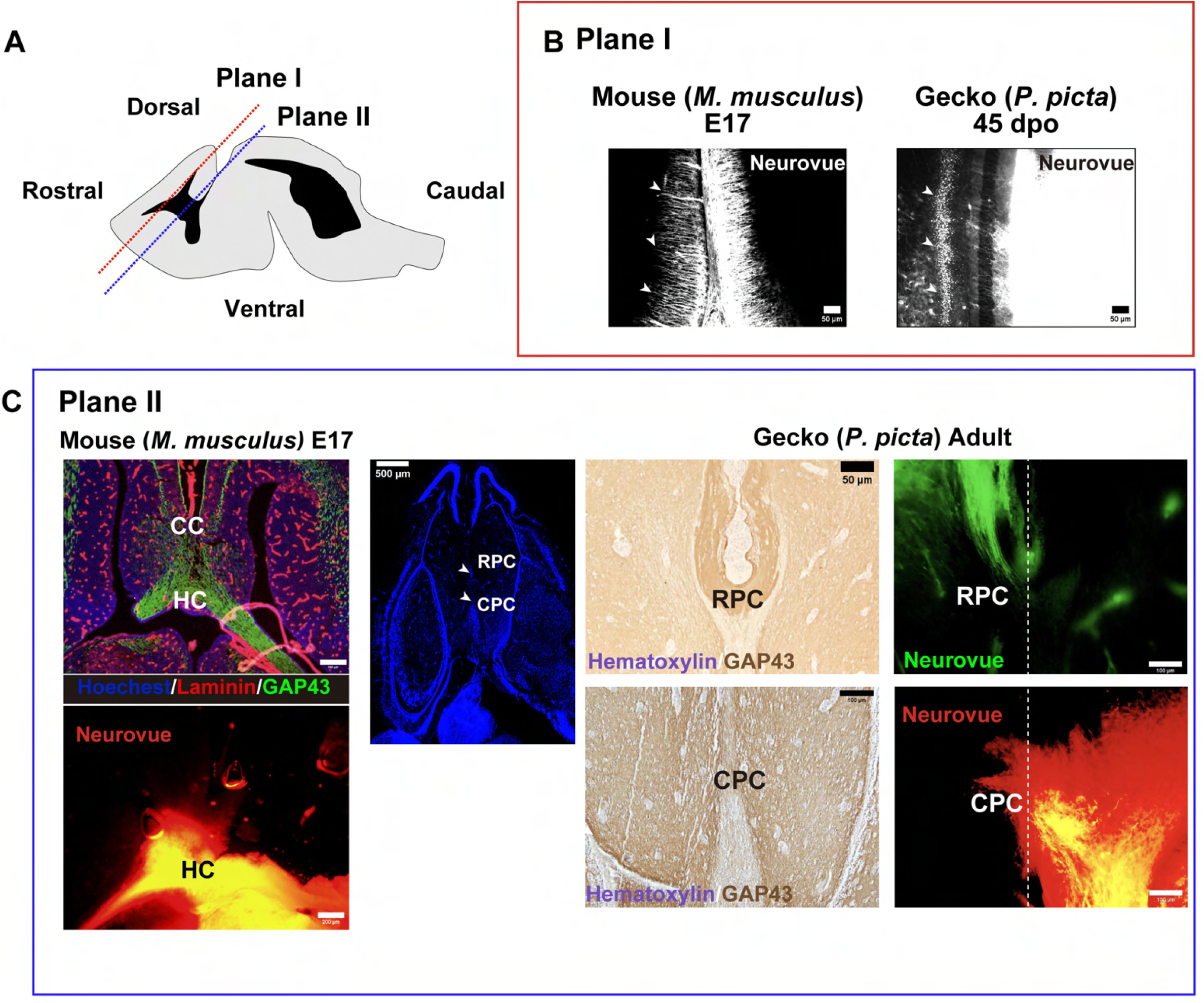
Rostral and caudal remodeling bridge the pathways of distinct pallial commissures. (A) A schematic drawing showing the position of plane I and plane II in the developing forebrain. (B) Retrograde tracing of callosal projection neurons in the mouse and pallial commissural neurons in the gecko (arrowheads in the left and right panels, respectively). The images correspond to plane I. (C) Immunohistochemistry and anterograde axon tracing showing the position of the corpus callosum (CC) and the hippocampal commissure (HC) in the mouse forebrain (left panels) and rostral and caudal pallial commissures (RPC and CPC, respectively) in the gecko forebrain (right panels).

### The absence of rostral IHF remodeling coincides with the lack of the RPC in chicks

Previously we reported that the minimal retraction of the IHF results in the absence of rostral interhemispheric remodeling in the developing chicken brain. To examine the route of pallial commissures that connect chicken telencephalic hemispheres, we performed anterograde axon tracing of distinct regions of the chicken pallium. The deposition of a lipophilic tracer on the hyperpallium apicale (HA), a homologous region to the mammalian neocortex, labeled axons that projected medially toward the rostral septum, whereas the tracer deposition on the hippocampus labeled discrete axonal projections direct to the caudal midline of the septum (Figure 5A-C). To further examine the projection patterns of HA-derived axons, we performed unilateral electroporation of GFP expression vector into the chicken brain. At E10, GFP-labeled axons derived from the HA extended ventrally toward the septum, routing to the ipsilateral ventral telencephalon via the tractus septomesencephalicus (TSM, Figure 5D). Importantly, these axons never crossed the midline of the septum in the developing chicken forebrain. Immunostaining with the GAP43 antibody also revealed ipsilateral axonal projections in the rostral septum (Figure 5D). To address whether the TSM is derived from specific neuronal subtypes, we performed retrograde axon tracing followed by immunostaining for Satb2, a transcription factor expressed in excitatory projection neurons. Injection of CTB488 into the rostral septum retrogradely labeled SATB2-positive neurons in the HA (Figure 5E). In developing chicken brains, caudal interhemispheric remodeling occurred at approximately E10 (Figure 1C). In contrast, the number and density of retrogradely labeled cells were not significantly different before (E9) or after (E13) remodeling. (Figure 5F), suggesting that caudal remodeling does not provide substrates for midline crossing of HA-derived axons. Using anterograde tracing and immunostaining with anti-GAP43 antibody, we confirmed that the CPC instead crossed the midline after caudal interhemispheric remodeling (Figure 5G, H). These data indicated unique trajectories of pallial axons in chicken hemispheres, and in line with the absence of rostral remodeling, axons derived from the HA do not cross the midline, unlike to mammalian CC and reptilian RPC.

**Figure 5.**
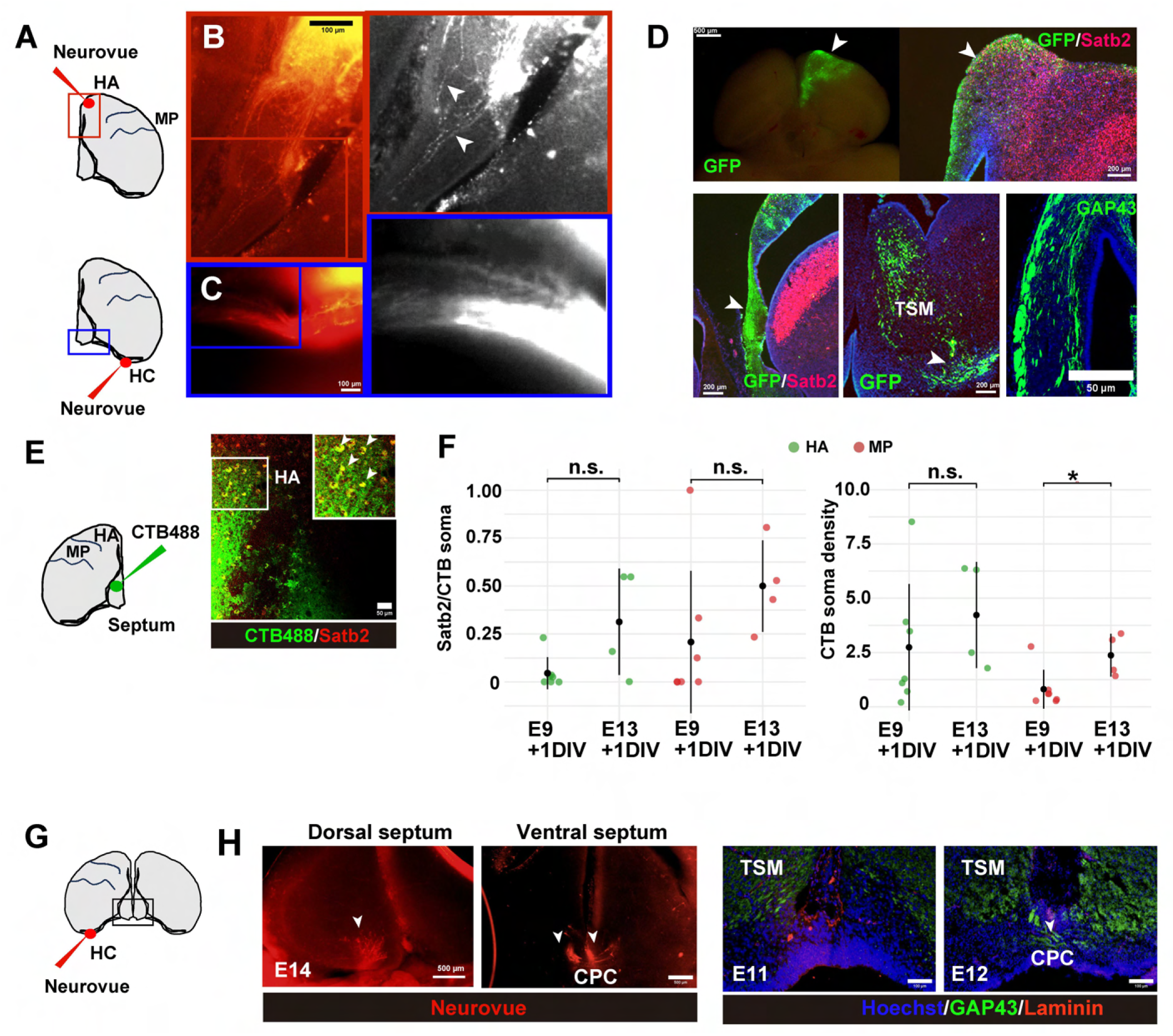
The absence of rostral IHF remodeling coincides with the lack of the RPC in the chick forebrain. (A) Schematic drawings showing injection sites of the lipophilic tracer in chicken brain sections. (B, C) Anterograde tracing of Hyperpallium apicale (HA)-derived axons in the rostral septum (B) and HC-derived axons crossing the caudal septum. (D) Anterograde tracing of HA-derived axons by electroporation of a GFP-expression vector Images were coronal sections. GFP-labeled axons projected toward the ipsilateral septum and contributed to the TSM. Image was coronal sections. Immunostaining with an anti-GAP43 antibody also revealed axons running through the septum Image was horizontal section. (E) Retrograde tracing of Satb2-positive neurons by injecting CTB488 into the septum. (F) Proportions of Satb2-positive soma in CTB-labeled soma (left panel) and CTB-labeled soma density (right panel) in chicken brain sections at different developmental stages. Soma located in HA or Mesopallium^54^ were labeled in green or red, respectively. E9+1DIV:N=7; E13+1DIV:N=4 (G) Anterograde tracing of the CPC at the caudal septal midline in the E14 chicken brain. GAP43-positive CPC was detected only the ventral septum where the IHF remodeling occurs.

### Genes for interhemispheric remodeling are expressed in the developing chicken brain

To further investigate the evolutionary conservation of gene expression associated with interhemispheric remodeling, we performed spatial gene expression profiling on the Visium platform (10x Genomics). We collected data from coronal sections of the forebrain hemisphere of a E14 chick. The gene expression of individual spots was automatically classified into 8 clusters and visualized by dimension reduction with *t*-distributed stochastic neighbor embedding (*t*-SNE, Figures S2A and S2B). Each cluster was mapped to a specific anatomical region in the chicken hemisphere (Figure S2A). We focused on cluster 8, which corresponded to the midline region as well as the margin of the hemisphere, presumably containing leptomeninges (Figure S2C). We found that several genes associated with midline glial components or interhemispheric remodeling, such as *Vimentin* (Figure S2D), *MMP2* (Figure S2E), and *Draxin* (Figure S2F), were also expressed in cluster 8, suggesting that molecular basis for caudal interhemispheric remodeling is highly conserved between mice and chicks, despite the lack of rostral remodeling and the RPC in chick brains.

### Satb2-dependent rostral remodeling is associated with CC and RPC formation in both mice and geckoes

Our comparative analyses suggest that discrete developmental mechanisms confer the rostral and caudal interhemispheric remodeling in amniotes. It has been shown that the rostral part of CC formation was severely compromised in cortex-specific *Satb2* mutant mice, in accordance with aberrant projection of callosal axons toward the ipsilateral septum^20^. Furthermore, several histological data from *Satb2* knockout mice implied incomplete IHF retraction in the rostral part of the septum^18,23^. These lines of evidence prompted us to re-examine the interhemispheric remodeling phenotype in *Satb2* mutants. To test this hypothesis, we generated *Satb2*-targeting mice using CRISPR-mediated gene editing (Figure S3A-S3C). Detailed histological examination revealed that callosal agenesis was closely associated with the disconnection of the rostral septal midline in the mice with only *Satb2* deletion alleles (*Satb2^del/del^*), whereas callosal formation and midline morphology were not affected in the mice with the wild-type allele (*Satb2^WT/del^*, Figure 6A). We detected altered accumulation of the GFAP-positive glial population including the MZG, indusium griseum glia, and glial wedge, on both sides of the midline in *Satb2^del/del^* mice (Figure 6B). Concomitantly, an altered direction of GFAP-positive glial fibers was evident in *Satb2^del/del^*mutants, suggesting that disorganized glial components contribute to the failure of IHF remodeling (Figure 6B, C).

**Figure 6.**
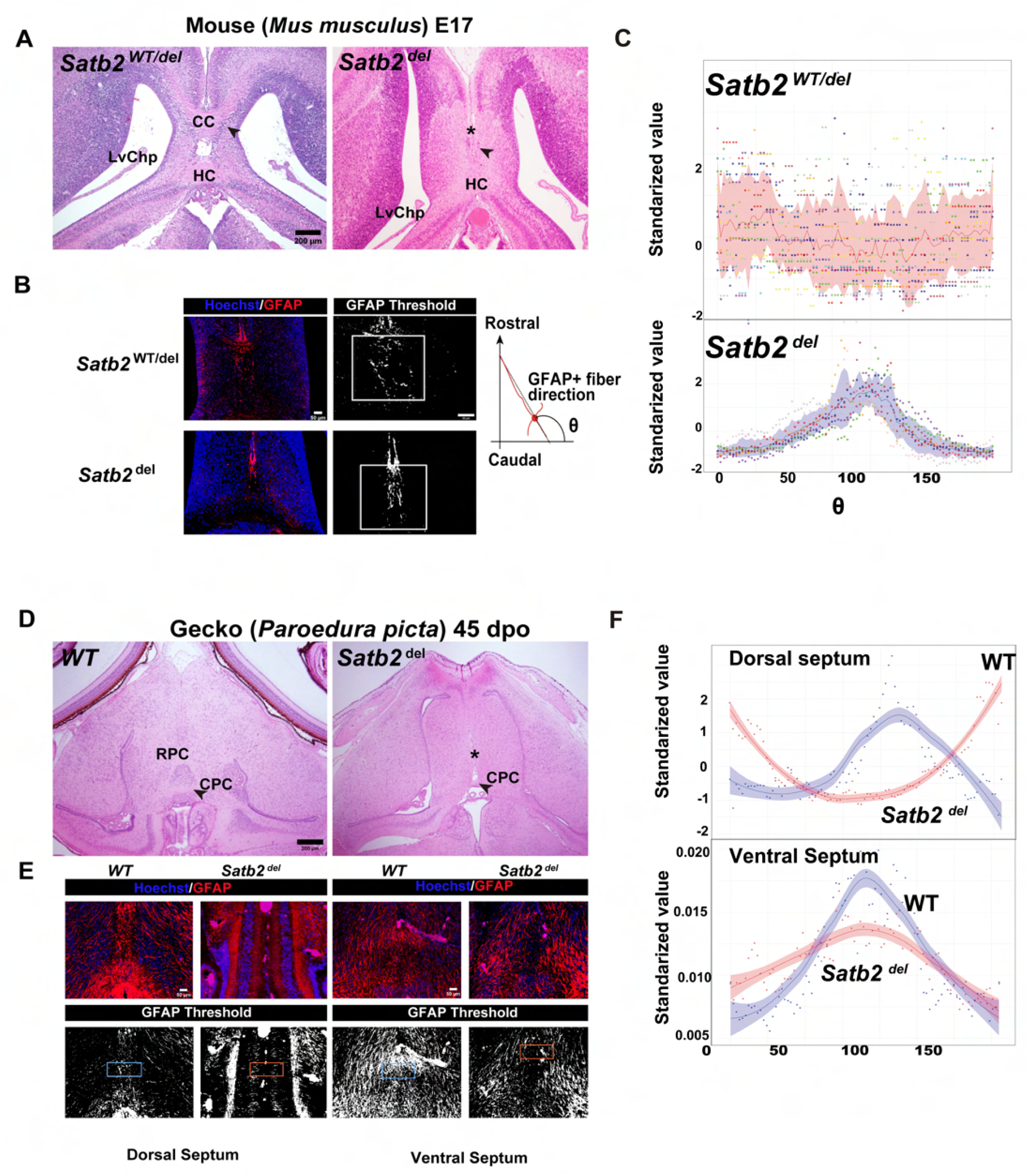
Satb2-dependent rostral remodeling is associated with CC and RPC formation in both mice and geckoes. (A) Horizontal sections of *Satb2^WT/del^* and *Satb2^del^*mouse forebrains. Arrowheads indicate the cellular accumulations corresponding to MZGs. (B) Distributions of GFAP-positive glial fibers in the *Satb2^WT/del^*(upper panels) and *Satb2^del^* (lower panels) IHF regions. Threshold processed images in higher magnification are indicated in black and white. White boxes indicate ROI (Region of Interest) for quantification. Schematic drawing showing how the direction of GFAP+ fiber is quantified. (C) Quantification of GFAP-positive fiber directions in the *Satb2^WT/del^*(upper panels) and *Satb2^del^* (lower panels) IHF regions. The different colors of each dot indicate different sections from the same individual. (D) Horizontal sections of wild-type (WT) and *Satb2^del^* gecko forebrains. An asterisk indicates the remanent of IHF. (E) Distributions of GFAP-positive glial fibers in the dorsal (left) and ventral (right) septum of WT and *Satb2^del^*geckos. Threshold processed images are indicated in black and white. Threshold processed images in higher magnification are indicated in black and white. Blue or red boxes indicate ROI (Region of Interest) for corresponding quantification. (F) Quantification of GFAP-positive fiber directions in the dorsal and ventral septum of WT and *Satb2^del^* geckoes. CC: Corpus Callosum. HC: Hippocampal Commissure. LVChP: Lateral ventricle Choroid Plexus.

Next, we investigated whether *Satb2* contributes to rostral interhemispheric remodeling and RPC formation in non-mammalian amniotes. We have previously shown that Satb2 is expressed in the developing gecko pallium^25,26^. Thus, we performed CRISPR-mediated gene targeting to disrupt *Satb2* in geckoes (Figure S4A, S4B). Compared with wild-type embryos, mutant gecko embryos with only *Satb2* deletion alleles (*Satb2^del/del^*) exhibited a shortened mandible (Figure S4C), which resembled Satb2-deficient mice^27^. Microcomputed tomography (micro-CT) analysis indicated abnormal skeletal formation and altered distribution of pan-collagen in the midline of *Satb2^del/del^* embryos (Figure S4C). Histological analysis revealed that *Satb2^del/del^*mutant embryos exhibited the disconnection of hemispheres and RPC agenesis (Figure 6D). The failure of IHF retraction was specific to the rostral septum, whereas the midline closure and CPC projection were not compromised at the caudal septum (Figure 6D). We examined whether the distribution of midline glial components was affected in the *Satb2^del/del^* mutant. In contrast to those in mice, GFAP-positive fibers extended in the rostral-caudal direction in wild-type gecko (Figure 6E, F). However, the vertical projection of glial fibers was less prominent in *Satb2^del/del^* geckoes (Figure 6E, F). These data suggest that the role of Satb2 in the rostral interhemispheric remodeling is highly conserved in eutherian mammals and reptiles, depending on species-specific distribution of midline glial cells in the dorso-ventral axis.

### Satb2 affects astrocyte characteristics in the developing mouse pallium

SATB2 is expressed in postmitotic neurons and plays an essential role in the specification of upper-layer neurons in the developing mouse neocortex^17,18,20^. Intriguingly, the loss of callosal projection identity also affects laminar-specific astrocyte identity in *Satb2* knockout mice^3^. To address the functional link between Satb2 and midline glial development, we performed gene expression profiling in E17.5 *Satb2* knockout mice based on published datasets^28^. The expression of callosal development-associated genes, such as *Epha4*^29^, *Ephb1*^29^, *Zeb2*^30,31^, *Robo2*^32^, and *Slit3*^15^, *Unc5C*^19^ were significantly altered in the cerebral cortex of *Satb2* mutant mice compared with than in wild-type mice (Figure S5A-C). To further investigate the contribution of Satb2 functions in glial cell characteristics, we secondary analyzed a previously published single-cell RNA-seq datasets of E15.5 mouse cortex^33^. We classified total 7 clusters (#0-6) with distinct cell states and identified lineage trajectories from cluster 4 toward cluster 1 and 3 (Figure S5D), which contained cells expressing genes associated with midline glial cells or interhemispheric fusions (Figure S5E, F). Intriguingly, *Satb2* was highly expressed in *Zeb2*-enriched cluster 4, and gradually decreased along with pseudotime (Figure S5D-F). By immunohistochemistry, we confirmed that SATB2 is expressed in the midline subventricular zone, adjacent to the glial wedge that produces GFAP-positive midline glial sling in the developing mouse brain (Figure S5G). Taken together, these results suggest that *Satb2* contributes to the establishment of midline glial cell characteristics.

## DISCUSSION

Here we demonstrate that interhemispheric remodeling, an initial step for CC formation in eutherian mammals, is also detected in non-mammalian amniotes. Spatio-temporal remodeling procedures coincide well with the routes of distinct pallial commissures, such as the RPC and CPC, in the reptilian forebrains. The reptilian and avian pallial commissures are considered to connect the hippocampus, therefore these axonal bundles are homologous to the HC^2,34,35^. However, several histological reports have suggested that the reptilian pallial commissure consists of the rostral and caudal pathways, and the former carries axons derived from the dorsal cortex, a homolog of the mammalian neocortex, whereas the latter connects the medial pallium, a homologue of the mammalian hippocampus^36^. In the present study we identified that the RPC and the CPC exhibit a high degree of similarity with the mammalian CC and HC, in terms of anatomical and developmental aspects. Thus, we propose that developmental mechanisms for interhemispheric connections are highly conserved in amniotes, and that the regulatory components for the RPC, probably derived from the ancestral amniotes, may correspond to evolutionary precursors for the development of the CC in eutherian mammals.

Our comparative analyses suggest that the rostral and caudal interhemispheric remodeling is conferred by distinct developmental mechanisms. In particular, caudal remodeling is mediated by the rearrangement of the lamina terminalis, leading to separation from the telencephalic hinge and a part of choroid plexus. The caudal remodeling occurs prior to midline crossing of the CPC or HC, suggesting that bridging mechanisms for the caudal septum midline provide substrates for the evolutionarily conserved commissural pathways. Notably, the rostral interhemispheric remodeling and subsequent formation of the RPC were not detected in chicken forebrain. Instead, the axons derived from the HA, a homologous region of the mammalian neocortex, extend to the ipsilateral septum and contribute to the TSM. We propose that avian TSM corresponds to the reptilian RPC, and that the rostral degeneration of IHF and midline crossing of RPC may have been secondarily lost during the evolution of avian lineages. Intriguingly, IHF degeneration is not detected in marsupials or monotremes that lack CC formation^11^. Whether these animal characteristics are the result of secondary modifications or are derived from ancestral traits awaits further analysis.

Disruption of Satb2 function results in impaired rostral interhemispheric remodeling in both mice and geckoes, suggesting evolutionary conservation of Satb2-dependent pallial rostral commissure development in amniotes. It has been considered that CC agenesis in *Satb2* mutant mice is due to misrouting of callosal axons to subcortical pathways, rather than impaired differentiation of midline glial cells^17,18,20,37^. In contrast, we demonstrated that in both mouse and gecko *Satb2* mutants, the failure of interhemispheric remodeling is associated with altered distribution of midline glial cells. Furthermore, the analysis of gene expression profiles in mice suggested that Satb2 plays a role in the establishment of astrocytic cell states. Because Satb2 is not expressed in differentiated midline glial components, we hypothesize that Satb2 has a non-cell autonomous effects on altered midline glial distribution. Alternatively, the defect in midline glial cells may be due to the secondary effects of impaired neuronal differentiation, as shown in a previous report of laminar-specific astrocytic differentiation^3^. Interhemispheric fissure is composed mainly of connective leptomeningeal fibroblasts and the extracellular matrix, which act as barriers to the callosal axon projection^12^. It has been reported that in *DRAXIN* or *Msx2-Ctnnb1* KO mutant mice, callosal agenesis is associated with pial membrane hyperplasia^8,13^. Several studies have reported that callosal abnormalities are linked with craniofacial abnormalities, suggesting a functional link between cranial mesenchymal differentiation and callosal projection. Intriguingly, *Satb2* is expressed in cranial mesenchymal cells of the osteoblast lineage^27^, thus it is possible that the failure of IHF retraction is secondary to cranial abnormalities.

Rostral interhemispheric remodeling does not occur in marsupials or monotremes; in these animals, axons from upper cortical neurons are directed to the anterior commissure^2^. Diffusion tensor imaging of human callosal dysgenesis suggested that the failure of midline crossing causes the rerouting of callosal fibers to various white matter regions^38^. A previous study indicated that species-specific routing of intracortical projections is mediated by differential timing of Satb2 expression^22^. Satb2 deficiency in mice has been shown to cause misrouting of callosal projections toward the ventral septum^20^, which partially resemble chick TSM routing. Although mechanisms underlying the developmental and evolutionary plasticity of axonal routing remain to be elucidated, our results suggest that Satb2 plays a key role in the multiple steps of pallial commissure development and evolution, which also provides a better understanding of variable human callosal agenesis.

## RESOURCE AVAILABILITY

## Lead Contact

Further information and requests for resources and reagents should be directed to and will be fulfilled by the Lead Contact, Tadashi Nomura.

## Material availability

All unique/stable reagents in this study are available from the Lead Contact with a completed Materials Transfer Agreement.

## Data and code availability

All original code and the data used to generate the figures and in this manuscript are have been deposited at Mendeley Data (DOI: 10.17632/2yx33zfkgn.3): The raw data generated in this study will be shared by the lead contact.

## ACKNOWLEDGMENTS

The authors thank Ms. Misato Kawami and Mariko Yazaki for their technical assistance, and Anna Yasunaga for providing wild type geckoes. This research was funded in whole, or in part, by Japanese Grant-In-Aid for Scientific Research (KAKENHI, 21H02594 to T.N.), the Takeda Science Foundation (to T.N.), the Ohsumi Frontier Science Foundation (to T.N.), Leading Initiative for Excellent Young Researchers (LEADER, 2020L0019) and JSPS KAKENHI-Grants (20K22665, 22H02638 to T.K.) and the Kyoto Prefectural University of Medicine 2121 Fellowship (to R.N.)., and Leave a Nest Co., 63rd incu·be encourage award (to R.N).

## AUTHOR CONTRIBUTIONS

Conceptualization, R.N. and T.N.; methodology, H.K., M.K., T.A, and Y.N.; investigation, R.N. and T.N.; visualization, R.N. and T.N. funding acquisition, R.N. and T.N. and T.K.; project administration, H.G.,C.O., K.O. and T.Y.; supervision, T.N.; writing – original draft, R.N.& T.N. writing – review & editing, all authors.

## STAR METHODS

The detailed methods are provided in the online version of this paper and include the following:

### KEY RESOURCES TABLE

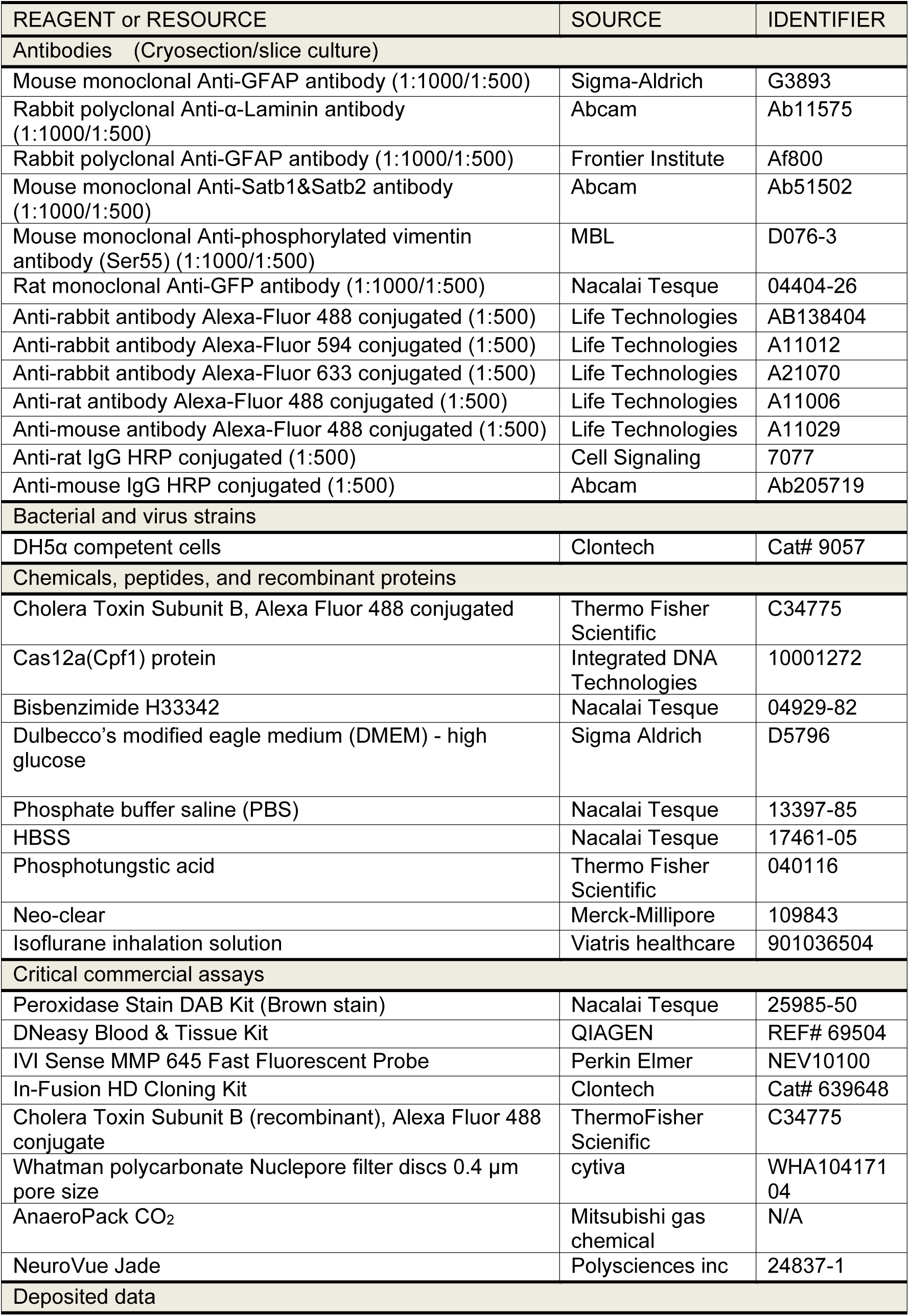
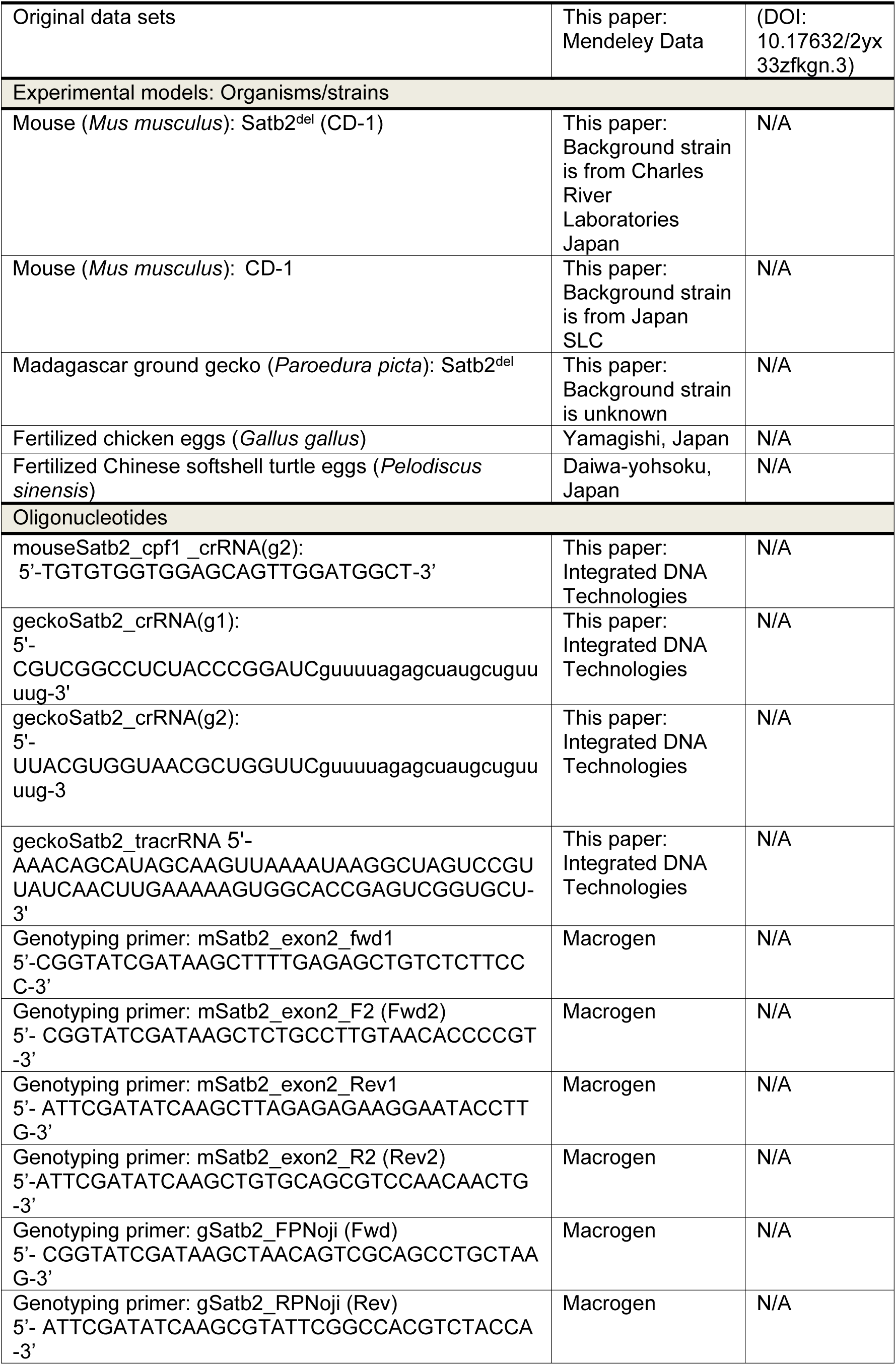
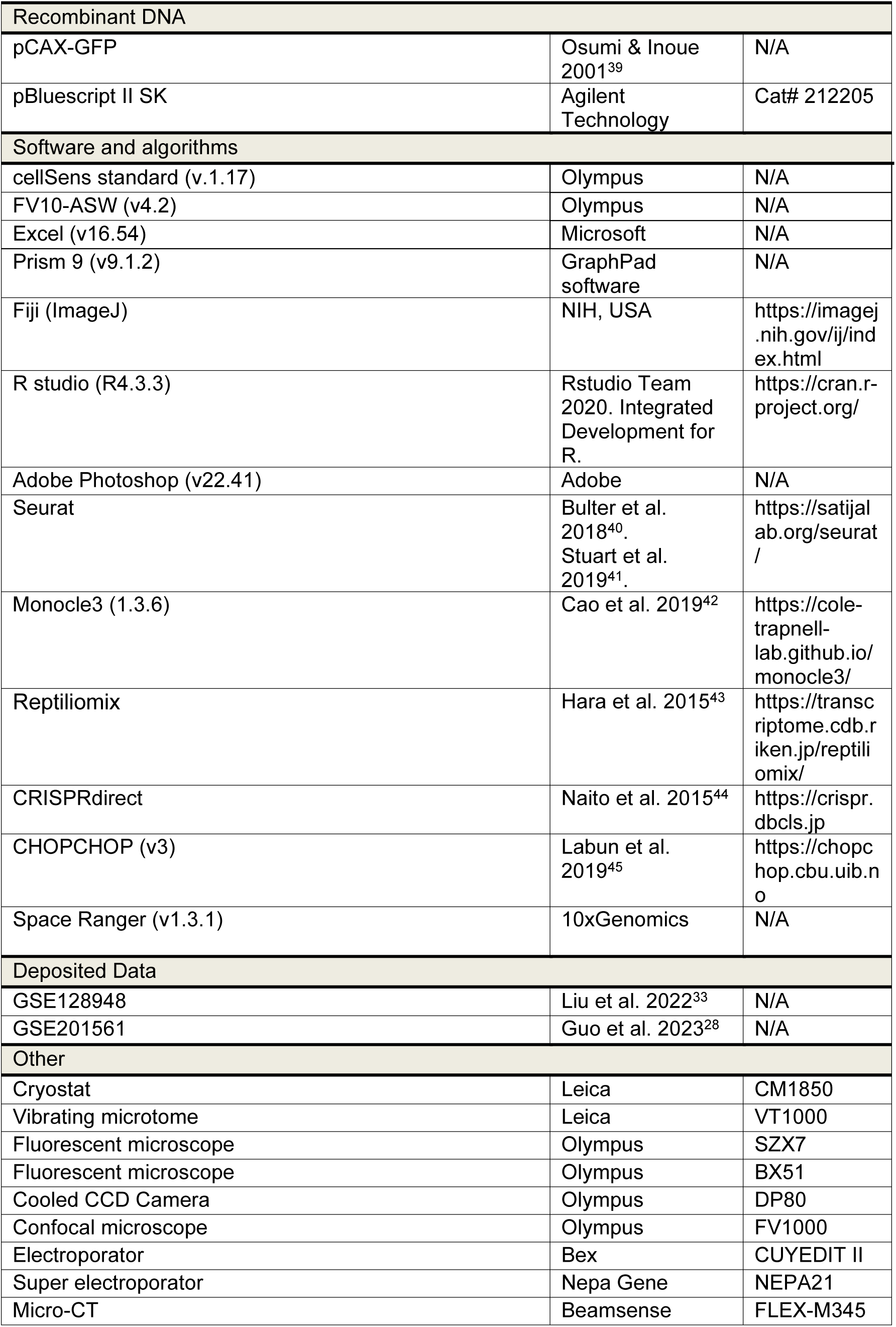

### METHODS DETAILS

#### Animals

Mouse embryos (CD-1 background) that were originally obtained from Japan SLC were maintained in at the experimental animal facilities of Kyoto Prefectural University of Medicine, Kyoto Institute of Technology in accordance with the relevant guidelines (M2023-199, M2023-202, M2023-205, R2023-5, #5), and the Institutional Animal Care and Use Committee of RIKEN Kobe Branch. Fertilized chicken eggs (*Gallus gallus*) were obtained from a local farm (Yamagishi, Japan) and incubated at 38 °C. The developmental stages of the embryos were assessed using a previously described method^46^. The fertilized Madagascar ground gecko eggs and adults were maintained in Developmental Neurobiology, Kyoto Prefectural University of Medicine and the animal facility of RIKEN BDR at 28 °C-30 °C until the analysis, and the embryonic stages were determined in accordance with previous studies^47^. Fertilized Chinese softshell turtle eggs were obtained from a local breeder (Daiwa Yoshoku) and incubated at 28 °C-30 °C until the time of analysis, and the embryonic stages of the turtles were determined according to a previous study^48^.

#### Paraffin sectioning

The embryonic samples were fixed with modified Clarke solution (99.5% ethanol and acetic acid at a ratio of 3:1). After rehydration, decalcification was performed using a 10% EDTA solution (pH 7.4) at room temperature for 1 week, followed by replacement with phosphate-buffered saline (PBS). The tissues were dehydrated using a series of ethanol solutions, and the samples were treated with Neo-Clear. Serial sections (4 µm thickness) were made with a microtome (Yamato Kohki) and mounted onto MAS coated glass slides (Matsunami) and incubated at 50 °C. After deparaffinization, the sections were subjected to hematoxylin and eosin (H&E) staining or immunohistochemical staining.

#### Immunohistochemistry

Embryonic samples were fixed overnight at 4 °C in a 4% paraformaldehyde (PFA)/1×PBS solution or a formalin-alcohol solution. After washing with PBS, cryoprotection was achieved with 20% sucrose solution, followed by embedded in OCT compound (Sakura, Japan). The frozen brains were sectioned at a thickness of 10 or 20 µm using via a cryostat. Alternatively, the fixed brains were sliced with a vibrating microtome. For immunohistochemistry, the sections were incubated with primary antibodies (see Key source table). After washing with Tris-buffered saline with Tween 20 (TBST) or PBS with Triton-X (PBST), the sections were incubated with secondary antibodies (see Key source table). Fluorescence images were captured with fluorescence microscopes equipped with a cooled CCD camera and a laser confocal microscope. All the captured images were processed with cellSens standard, FV10-ASW, ImageJ and Adobe Photoshop.

#### *Ex vivo* slice culture

Embryonic chick or turtle brains were dissected in 1×PBS and embedded in 4% low-melting agarose for approximately 10 min in room temperature. To prevent autofluorescence, chicken embryos were perfused with ice-cold PBS before isolation of brains. Embedded brains were sliced with a vibrating microtome with 250 or 300 µm in thickness. Brain slices were placed on a culture filter and cultured in DMEM plus 10% FBS and 1% antibiotics (penicillin/streptomycin). The detection of pan-matrix metalloproteinase (MMP) activity in the developing Chinese softshell turtles was performed according to a previous report^12^ with slight modifications. Briefly, brains slices were incubated the culture medium containing 0.4 µM MMPSense 645 Fast fluorescent probe at 30 °C or 37 °C for 24 hours.

#### Axon tracing

After the brain was fixed with 4% PFA/1×PBS, small fragments of filters containing lipophilic dye (NeuroVue) were inserted into the specific brain regions. The samples were incubated in 1% PFA/1×PBS at 37 °C for 3–4 weeks, then analyzed following sectioning with a vibrating microtome. Alternatively, the fixed brains were sectioned with a vibrating microtome prior to the axon tracing. For retrograde axon tracing, isolated brains were sliced with a vibrating microtome, and Cholera Toxin Subunit B conjugated with Alexa fluor 488 (CTB488) was injected into the focal region of the slice. After injection, the slices were placed on a culture filter and cultured for 24 hours in 37 °C.

#### Generation of *Satb2*-deficient mice

*Satb2*-deficient mice were generated via CRISPR/Cas12a (Cpf1)-mediated genome editing. A crRNA sequence that targets mouse *Satb2* exon 2 was designed by CHOPCHOP. We chose a target sequence with no off targets on the basis of mouse genome information (mm10/GRCm38). Genome-edited mice on the CD-1 background were generated via the *i*-GONAD method^49^. Briefly, pregnant female mice (E0.75) were anesthetized with 2% isoflurane and the CRISPR/Cas12a solution was injected into the oviduct lumen. Then, square electric pulses were applied to the oviduct via a pulse generator (Nepa gene).

#### Generation of *Satb2*-deficient geckoes

*Satb2*-deficient geckoes were generated via CRISPR/Cas9-mediated genome editing as previously described^50^. The predicted genomic sequences and cDNA of gecko (*Parodedura picta*) *Satb2* were identified by the Reptiliomix BLAST search (https://transcriptome.cdb.riken.jp/reptiliomix/). The predicted motif of the gecko Satb2 coding region was identified by MOTIF search (https://www.genome.jp/tools/motif/). gRNA sites were designed via CRISPRdirect (https://crispr.dbcls.jp) and specificity was checked based on Ppicta_v2.0. The female geckoes were anesthetized with an isoflurane inhalation solution. Microinjection into the female ovary was performed using an electric microinjector system (IM-400, NARISHIGE) under a stereomicroscope (Leica MZ95). After microinjection, the ovary was returned to the abdominal cavity and the musculature and skin were sutured.

#### Genotyping of *Satb2*-deficient mice and geckoes

Genomic DNA from embryonic mouse or gecko tissues (skin or tail fragments) was isolated via a DNeasy Blood & Tissue kit. The DNA fragments containing the CRISPR target region were amplified via polymerase chain reaction (PCR). To determine of the edited sequences in F0 embryos, PCR amplicons were subcloned into the pBluescriptSK vector, and six to ten randomly selected clones were examined via Sanger sequencing.

#### Electroporation

*In ovo* electroporation of developing chick embryos was performed according to a previous study^25^. Briefly, a window was opened in the shell of an egg, and then a small amount of DNA solution (less than 0.05 µL) was injected into Hamburger & Hamilton ^51^ 25 chicken embryos with a glass needle. Next, needle-type electrodes (CUY200S, BEX) were placed on the neural tube and square electric pulses (28 V for 50 msec, 3 times) were applied with an electroporator. After electroporation, the extraembryonic cavity was filled with sterilized Hank’s balanced salt solution (HBSS) containing antibiotics (1:1000; penicillin/streptomycin, gentamicin), and the window was sealed with cellophane tape. The electroporated embryos were incubated in an incubator at 38 °C Until the prescribed developmental stage.

*In utero* electroporation was performed according to previously described methods^52^. Briefly, a pregnant mouse was anesthetized with isoflurane, and an incision was made in the abdominal wall to expose the uterine horns. A small amount of DNA mixture (0.3– 0.5 µL) was injected into the lateral ventricle of each embryo, and square pulses (35 V, 50 ms, four times) were applied to the embryos with a tweezer-type electrode (CUY650P3, BEX) connected to a pulse generator (BEX).

#### IHF measurement

IHF measurements on H&E-stained samples were performed by using Fiji (ImageJ). IHF lengths were normalized to the entire rostral–caudal length of the terminal brain along the interhemispheric midline as previously described^12^. To determine macroscopic trends, a single regression line was drawn by using the least-squares method, and the direction and degree of remodeling were quasi-investigated by the slope. In this case, a single regression line was obtained and drawn for each developmental stage group of mice and geckos, respectively, via R software (ggplot2). The confidence intervals were set at 95%, and the ranges are shaded in the corresponding figure colors.

#### Spatial transcriptome

The brains of the embryos were harvested at chick E14, and directly embedded with O.C.T. compound. Embedded brain stored at −80 °C deep freezer until sectioning. Frozen sections were prepared from each brain at a thickness of 10 μm as manufacture’s protocol. The sections were placed on the Visium Spatial Gene Expression Slide (10x Genomics, USA) and processed with the Visium Spatial Gene Expression Reagent kit (10x Genomics, USA) according to the manufacture’s protocol. H&E brightfield and FL staining images were collected as mosaics using a Keyence BX-800 microscope (Keyence, Japan). The resultant libraries were sequenced at Macrogen inc. by the HiSeqX platform (Illumina, USA) with 150bp paired end sequencing.

#### Data processing of spatial gene expression

FASTQ files were processed with Space Ranger 1.3.1 with default parameter settings referring to the gene annotations of the galGal6 genome assembly provided by UCSC (galGal6.ncbiRefSeq.gtf.gz) resulting in a production of the matrices of gene expression profiles of the individual cells. Its FASTQ files were retrieved from the UCSC genome browser(https://hgdownload.soe.ucsc.edu/goldenPath/galGal6/bigZips/galGal6.fa.gz). The gene expression profiles of the E14 sample were processed with Space Ranger. Read count matrices were normalized, and the spots of the sample were grouped based on their expression profiles by k-means clustering, resulting in 8 clusters. 2D *t*-SNE plots were visualized with Loupe Browser application. Marker genes, those with significantly higher expression levels in a cluster than the others, were detected with Loupe Browser. The clusters were manually annotated referring to the ‘known’ marker genes that were included in those automatically inferred.

#### RNA-seq data analysis

Bulk RNA sequencing data from the cortex of E17.5 wild-type and Satb2 KO mice were downloaded from GSE201561^28^. The tsv files were converted to csv files on R studio. DEGs of genes expressed in parts of the glial sling in the literature described in the text were compared via hierarchical clustering and violin plots in wild-type and *Satb2* KO mutant. Single-cell RNA-sequencing data of the E15.5 mouse astrocyte progenitor cells (GFP-positive and RFP-negative cells derived from Nes-GFP/Dcx-mRFP double transgenic reporter mice.) were downloaded from GSE128948. We used the Seurat package (v5.0.3) to analyze the data. Dimensionality reduction was performed via principal component analysis (PCA) and UMAP via the Seurat pipeline with standard parameters. R studio (package: SeuratWrapper, tidyverse, dplyr, SeuratWrappers, ggplot2, ggridges, and Monocle3) were used to draw violin plots and UMAPs, and Monocle 3 was used for psedotrajectory analysis.

#### X-ray computed tomography

For micro-CT scanning analysis, samples fixed with 4% PFA in PBS were stained with 1% phosphotungstic acid solution (w/v) in 100% EtOH and scanned with a FLEX-M345 at the Kyoto Institute of Technology. The scanned data were processed with Kyoto CT OD reconstruction or the original version of the software and were converted to a stack of 8-bit tiffs to hyperstack with Fiji (ImageJ). A peak voltage of 40 kV and a current of 100 µA, unfiltered, were used for skeletal staining data, and a peak voltage of 40 kV and a current of 80 µA were used for pan-collagen visualization data.^53^ For quantification of collagen in the central and middle of the range of equivalent anatomical positions, data were converted at 8 bits and the ImageJ plug-in (Plot Z axis) was used. To accurately reflect x-ray absorbance, the change in the same region of interest (ROI) over the dorsoventral axis of the embedding material-only region was subtracted and the values were standardized for PCA as described above. For statistical analysis, at least three independent samples from each experimental group were compared. Comparisons between experimental groups were performed using Microsoft Excel (v16.54, Microsoft) and R studio (package: ggplot2).

#### Image processing and quantificational analyses

All fluorescence images were captured with a fluorescence microscope equipped with a CCD camera (DP80, Olympus), and a confocal microscope. Images were examined and analyzed by using Fiji ImageJ. All images were processed with ImageJ and Adobe Photoshop 2024. Fluorescent images were displayed in pseudo-colour, which differs from the original colour if necessary. Glial fiber orientation was compared using Fiji/ImageJ software. Regions of a telencephalic hemisphere showing distinct fiber structures were extracted from paraffin or frozen sections of spoonbills and geckos stained with anti-GFAP antibody. The orientation pattern of the glial fibers in each image was calculated as a ‘Fourier component’ using the ‘Directionality plugin’ (http://fiji.sc/wiki/index.php/Directionality). The slice images used for analysis were adjusted using the auto threshold to select Max entropy of the Fiji plugin. The resulting csv file was loaded into R Studio and z-score normalization to the number of GFAP-positive fibers in a given direction was performed to examine the differences in the distribution map of GFAP-positive fibers. The standardized values output by the Directionality plugin were approximated using the R library (ggdist, dplyr, patchwork) and Gaussian fitted at the 95% confidence interval, and the ranges are shaded in the corresponding figure colors.

#### Data visualization and statistical analysis

To visualize the differential expression of genes, we utilized the ggplot2 package in R. We first imported the gene expression dataset using the read.csv and specified the file path of the data. To focus on key genes involved in interhemispheric remodeling we filtered the data, extracting rows corresponding to these gene symbols for further analysis. Next, we created a custom color-coding system based on the basis of log_2_-fold changes and adjusted *p-values*. Specifically, data points with a log_2_ fold change greater than 1 and an adjusted *p-value* less than 0.05 were colored red, indicating significant upregulation. Conversely, data points with a log_2_ fold change less than −1 and an adjusted p-value less than 0.05 are colored blue, indicating significant downregulation. Data points not meeting these criteria were excluded from the visualization by filtering out missing (NA) values in the color column. A volcano plot was generated using ggplot2, with the log_2_ fold change on the x-axis and the negative log_10_ value of the adjusted p-value on the y-axis. This plot allowed for the clear identification of significantly upregulated and downregulated genes. We enhanced the visualization by adding a horizontal dashed line to indicate the false discovery rate (FDR) threshold of 0.05, aiding in the interpretation of statistical significance.

CTB-labeled soma density was measured on confocal images of the four experimental groups using ImageJ. The data for each group were transformed by multiplying by 10^-4^ µm^2^ for consistency. For the analysis of neuronal soma density, we used ggplot2, dplyr, and ggpubr in the R environment. Given the presence of missing values in certain groups, tests were performed only on the available data points. On the basis of the results of the normality test, the appropriate statistical test was selected. If both groups passed the normality test (*p-value*>0.05), a *t*-test was performed to compare the means. Otherwise, a nonparametric Wilcoxon rank sum test was performed.10^-4^ µm^2^ for consistency. This approach was applied for both the comparisons, with corresponding p-values recorded for each test. The mean values and SEM (standard error of the mean) were added for each group via a point-range plot to provide visual context for the overall distribution of the data.

#### Data availability

The raw data of spatial transcriptomics have been deposited in the DDBJ data base (DRA BioProject: PRJDB15113). All data generated in this study have been deposited in Mendeley Data (Mendeley Data, V3, DOI: 10.17632/2yx33zfkgn.3).

