## Supplementary information for "Conserved interhemispheric morphogenesis in amniotes preceded the evolution of the corpus callosum"

**Figure S1. Distributions of GAP43-positive axons during IHF remodeling in the developing chick and turtle (related to Figure 4).**

(A-D) GAP43-positive axon bundles detected on both sides of the septum at E7 (A), E10 dorsal septum (B), and E10 ventral septum (C) in the developing chicken forebrains. GAP43-labeled axon bundles were also detected on both sides of the septum in the developing turtle forebrain (St. 21).

**Figure S2. Expression levels of genes associated with midline glial cells according to spatial transcriptomics (related to Figure 5).**

(A) A schematic drawing of the chicken forebrain. (B) Cluster identification of genes expressed in E14 chick forebrain. *t*-SNE visualization based on the transcriptional signatures of each spot. (C) Gene expression in cluster 8. (D-F) Expression of midline glial marker genes including Vimentin (D), MMP2 (E), and Draxin (F) in the forebrain (left panels). *t*-SNE visualization (center column), and expression levels of each gene in distinct clusters (right panel).

**Figure S3. Generation and characterization of Satb2-deficient mice (related to Figure 6).**

(A) A schematic drawing of the mouse *Satb2* genomic structure (upper panel). A guide RNA was designed to target the N-terminus of the Satb2 protein (lower panel). (B) The result of Sanger sequencing results for each crispant embryo. The blue color shows the PAM sequence, and red color indicate target sequences of the guide RNA. (C) A smaller jaw was evident in $Satb2^{del}$ mouse compared to $Satb2^{WT/del}$ mouse.

**Figure S4. Generation and characterization of Satb2-deficient geckoes (related to Figure 6).**

(A) A schematic drawing of the Madagascar ground gecko *Satb2* genomic structure (upper panel). A guide RNA was designed to target the C-terminus homeodomain (HD) of the Satb2 protein (lower panel). (B) The result of Sanger sequencing results for each crispant embryo. The blue color shows the PAM sequence, and the ed color indicates the target sequences of the guide RNA. (C) A smaller jaw was evident in $Satb2^{del}$ gecko than in WT gecko (left panels). Micro-CT analysis indicated anomalies of premaxillary (pmx) and mandibular bones (mb) in $Satb2^{del}$ gecko. Micro-CT images with 1% PTA staining and subsequent PCA analysis suggested altered collagen distribution in a dorsal-ventral axis in $Satb2^{del}$ geckoes, compared with WT geckoes. Mx: maxillary bone.

**Figure S5. Satb2 affects astrocyte characteristics in the developing mouse pallium, related to Figure 6**

(A) Volcano plot illustrating differentially expressed genes (DEGs) in the Bulk RNA-sequencing data of *Satb2* KO and WT neocortex[27]. (B) Heatmap visualization showing the expression of callosal development-associated marker genes in KO and WT samples. (C) Expression levels of astrocyte marker genes in Satb2 KO and WT samples. (D) U-MAP visualization of single-cell RNA-sequencing data from the developing mouse E15.5 astrocyte progenitors[31]. Different colors indicate distinct clusters of cells. Pseudotime trajectory analysis suggests lineage relationships of cell clusters (right panel). Satb2 expression was detected in cluster 4 (lower panels). (E, F) Expression of midline glial sling-associated genes in each cluster. (G) Lineage tracing (E14.5 to E16.5) of midline glial cells by *in utero* electroporation (IUE) of a GFP-expressing vector.SATB2 is expressed in subventricular progenitors as well as cortical neurons, which are close to the glial sling-derived MZG cells.

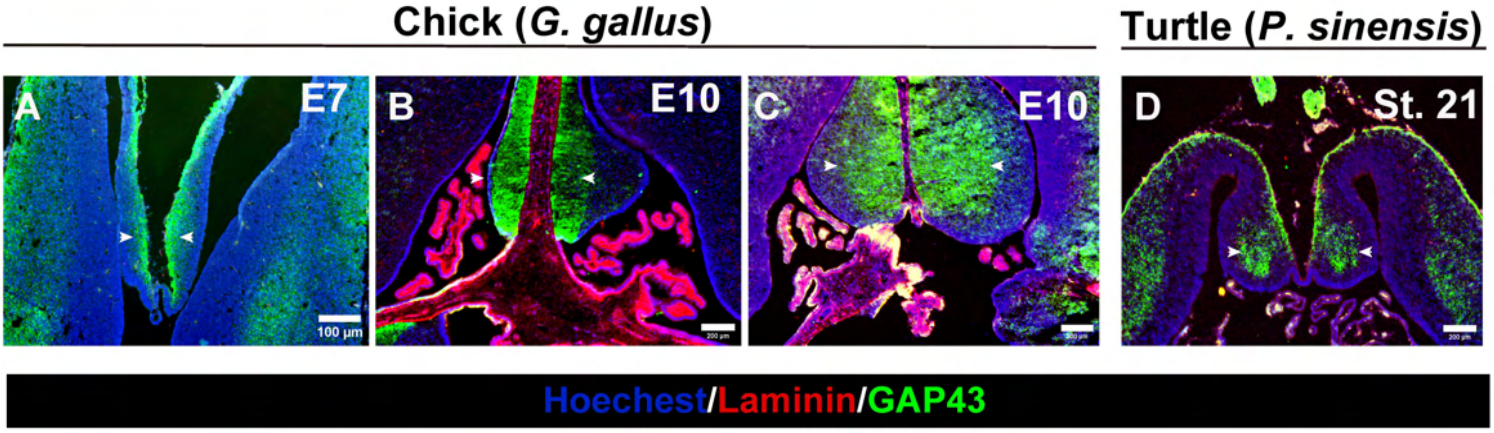


Noji et al. Figure S1

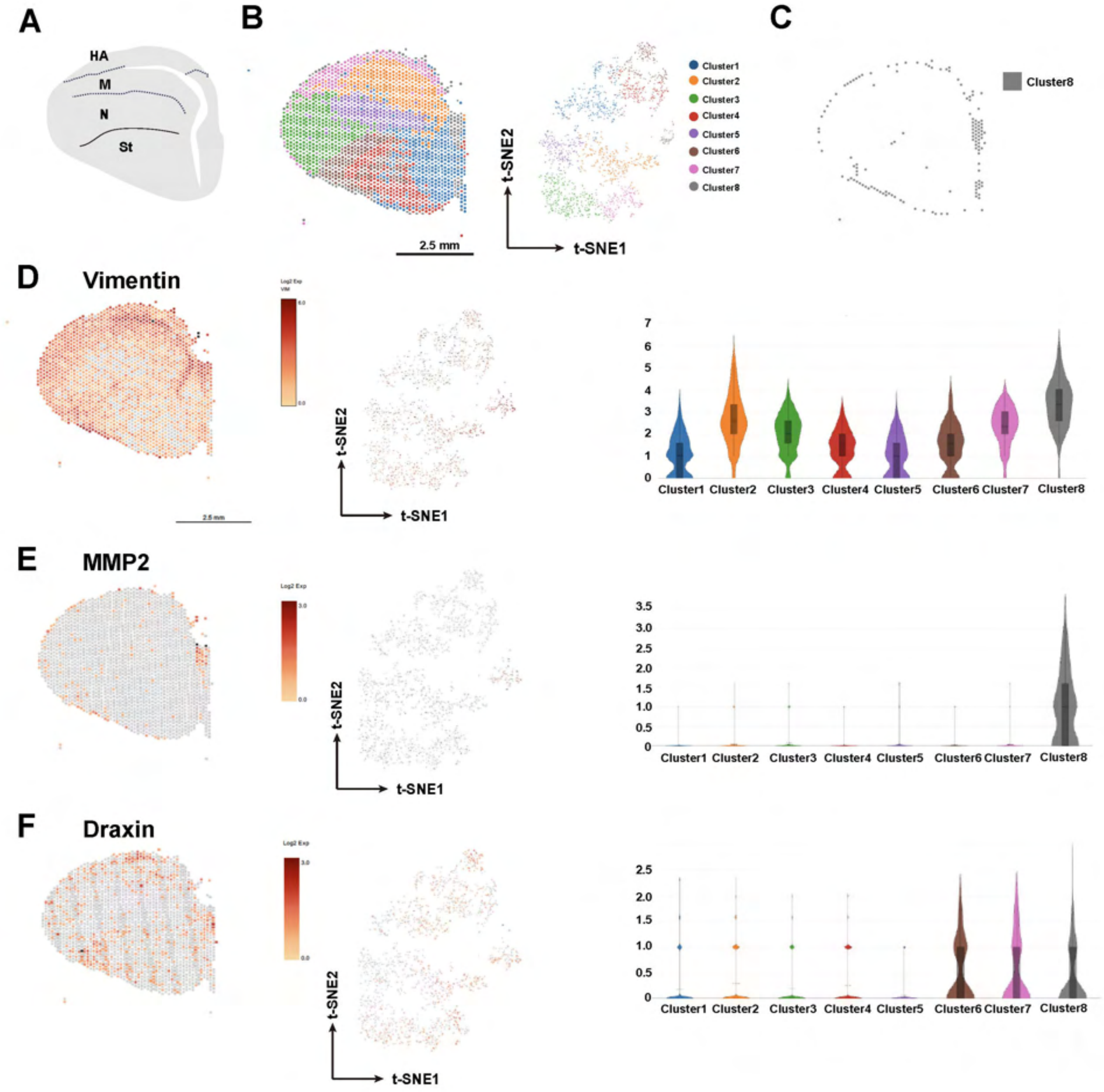


**Noji et al. Figure S2**

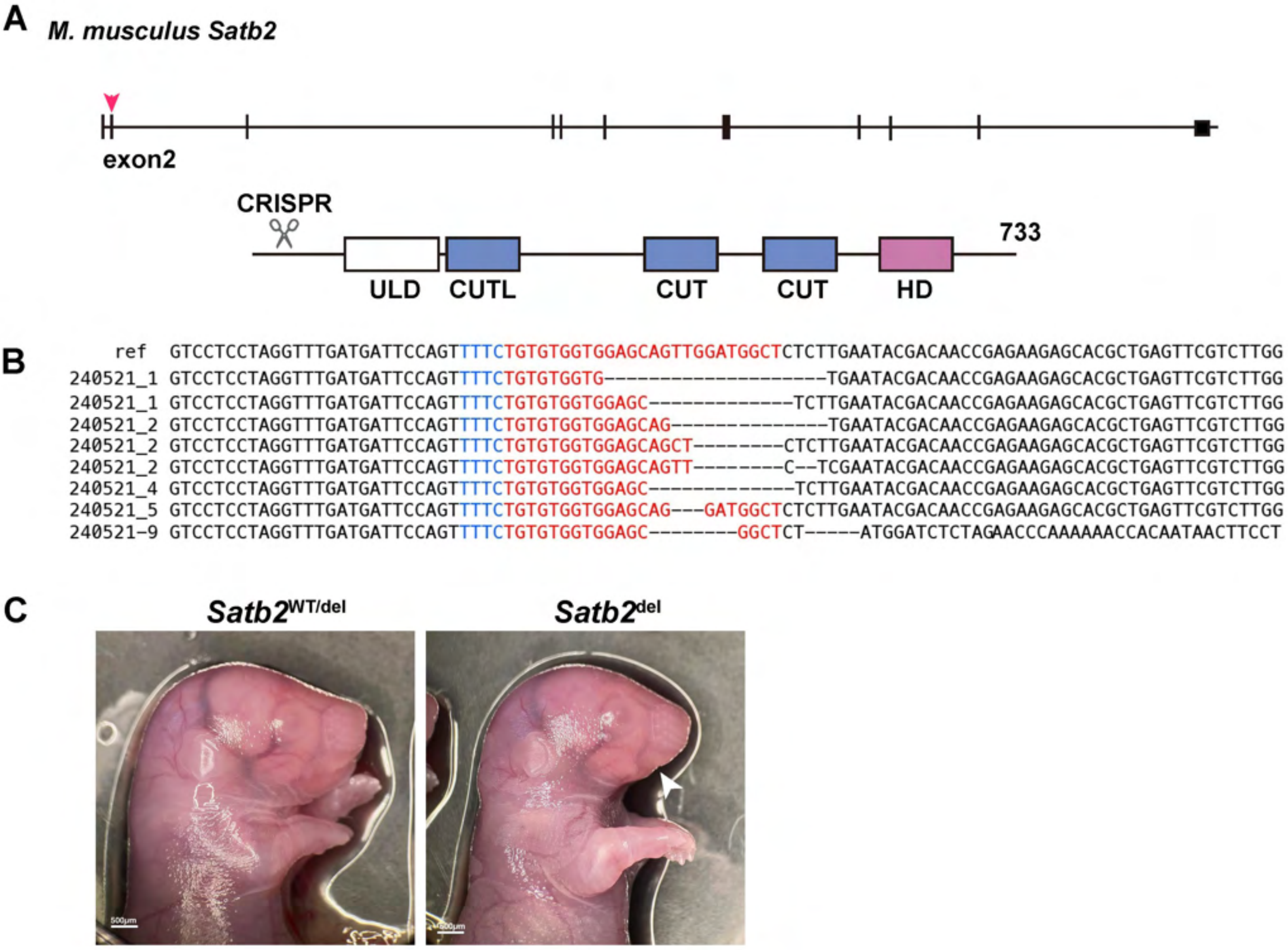


**Noji et al. Figure S3**

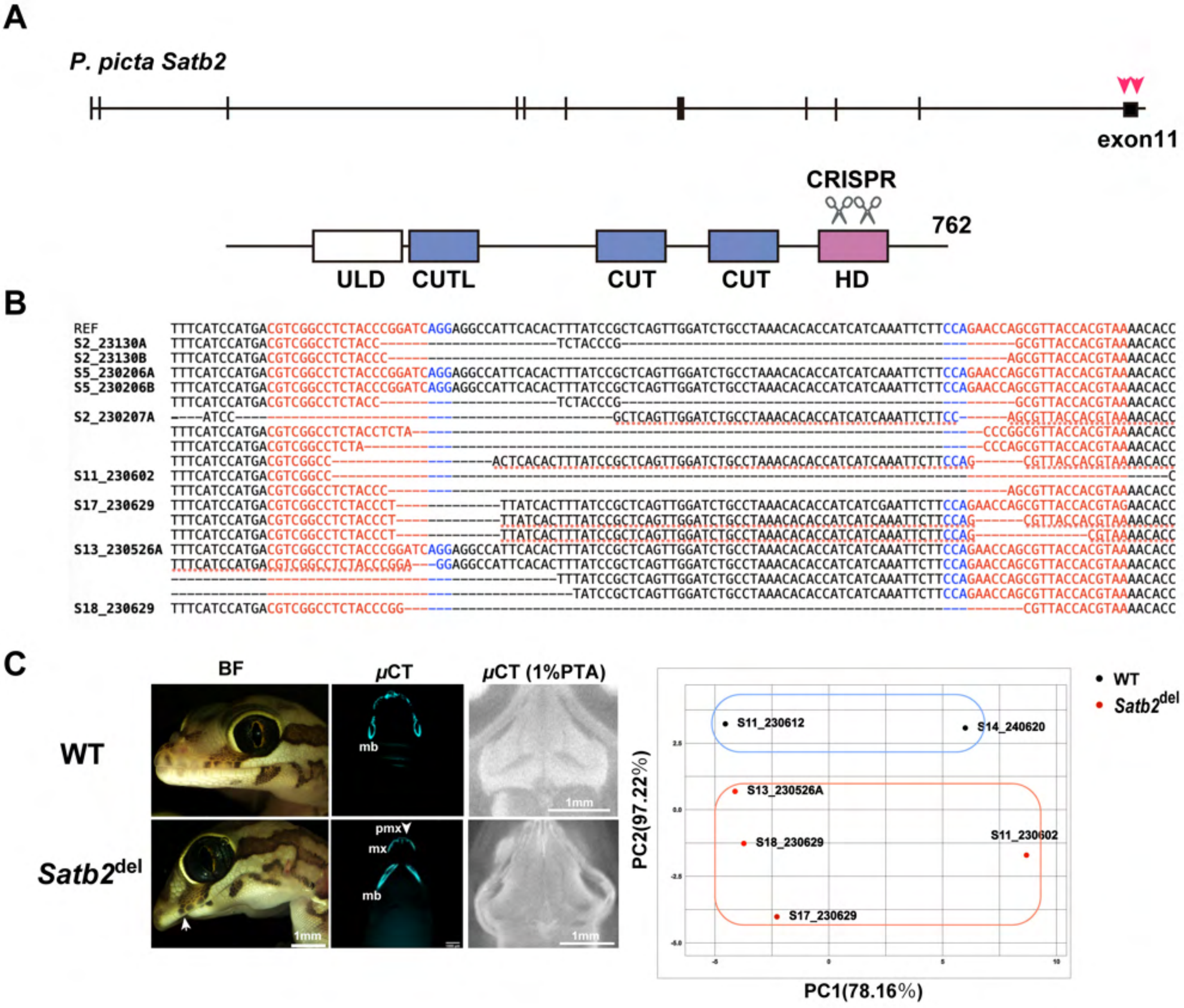


Noji et al. Figure S4

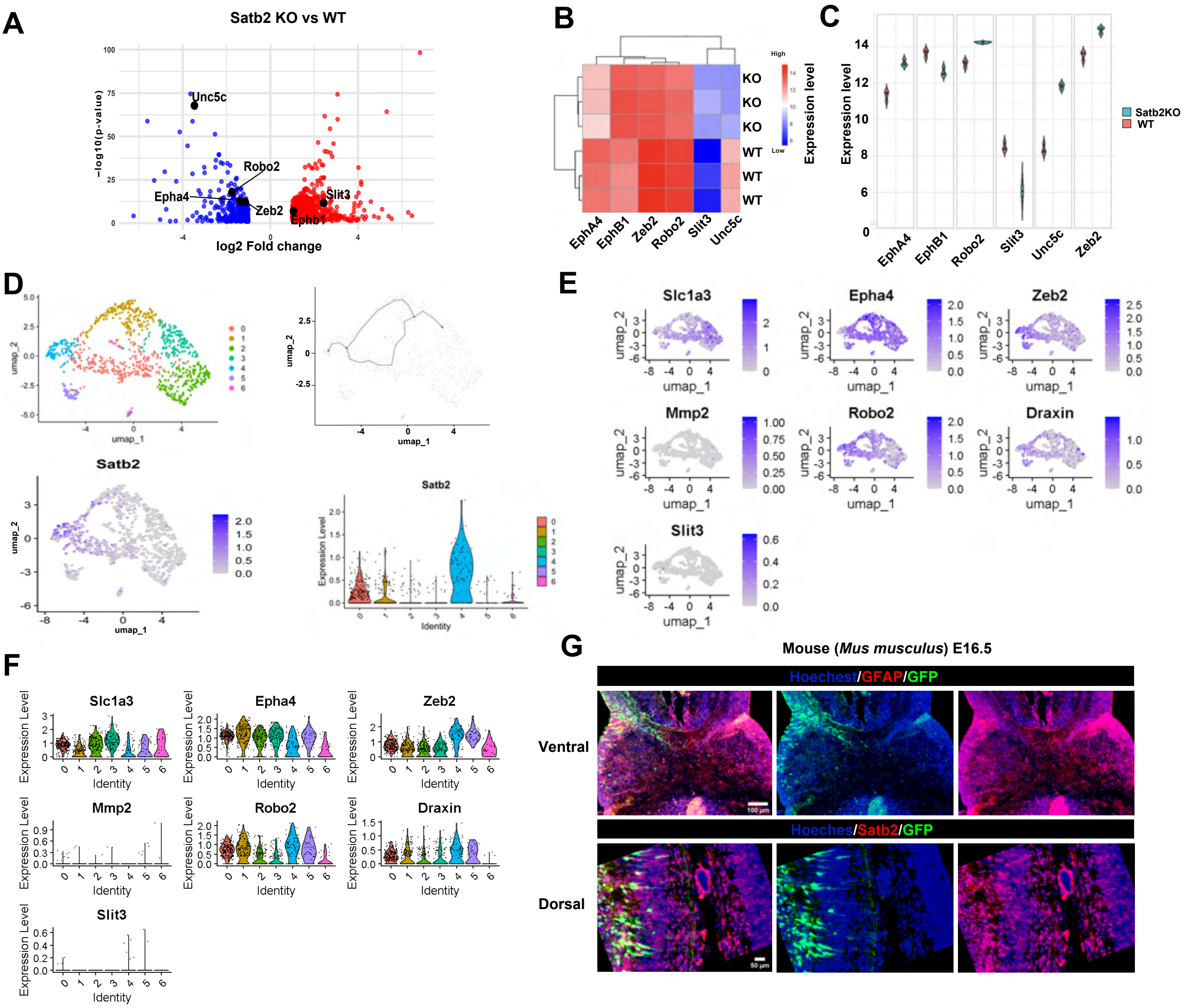


### Noji et al. Figure S5
